# Caveolin scaffolding domain (CSD) peptide LTI-2355 modulates the phagocytic and synthetic activity of lung derived myeloid cells in Idiopathic Pulmonary Fibrosis (IPF) and Post-acute sequelae of COVID-fibrosis (PASC-F)

**DOI:** 10.1101/2023.12.01.569608

**Authors:** Brecht Creyns, BreAnne MacKenzie, Yago Sa, Ana Lucia Coelho, Dale Christensen, Tanyalak Parimon, Brian Windsor, Cory M. Hogaboam

## Abstract

**Rationale:** The role of the innate immune system in Idiopathic Pulmonary Fibrosis (IPF) remains poorly understood. However, a functional myeloid compartment is required to remove dying cells and cellular debris, and to mediate innate immune responses against pathogens. Aberrant macrophage activity has been described in patients with Post-acute sequelae of COVID fibrosis (PASC-F). Therefore, we examined the functional and synthetic properties of myeloid cells isolated from normal donor lung and lung explant tissue from both IPF and PASC-F patients and explored the effect of LTI-2355, a Caveolin Scaffolding Domain (CSD) peptide, on these cells.

**Methods & Results:** CD45^+^ myeloid cells isolated from lung explant tissue from IPF and PASC-F patients exhibited an impaired capacity to clear autologous dead cells and cellular debris. Uptake of pathogen-coated bioparticles was impaired in myeloid cells from both fibrotic patient groups independent of type of pathogen highlighting a cell intrinsic functional impairment. LTI-2355 improved the phagocytic activity of both IPF and PASC-F myeloid cells, and this improvement was paired with decreased pro-inflammatory and pro-fibrotic synthetic activity. LTI-2355 was also shown to primarily target CD206-expressing IPF and PASC-F myeloid cells.

**Conclusions:** Primary myeloid cells from IPF and PASC-F patients exhibit dysfunctional phagocytic and synthetic properties that are reversed by LTI-2355. Thus, these studies highlight an additional mechanism of action of a CSD peptide in the treatment of IPF and progressive fibrotic lung disease.

## Introduction

IPF is an aging associated progressive fibrotic lung disease hallmarked by aberrant fibroblast and epithelial cell activity. The role of the immune system in the initiation and maintenance of this disease remains poorly understood. Myeloid cells including macrophages and dendritic cells are major innate immune cells, which play a central role in lung homeostasis, response to infection or injury, and tissue repair (Byrne et al., 2016). Lung-resident alveolar macrophages (TR-AMs) are the main AMs located, whereas in IPF ontologically distinct AM derived from circulating monocytes (Mo-AMs) drive disease (Misharin et al., 2017). More recently, single cell RNA sequencing (scRNAseq) efforts identified activated myeloid cell subpopulations that contribute to the fibrotic response (Adams et al., 2020; Morse et al., 2019; Reyfman et al., 2019). These scRNAseq analyses revealed at least 4 subpopulations based upon unique transcript expression including an osteopontin (SPP1)^+^ and Chitinase 3-like-1 (CHI3L1) high(^hi^) AMs specific to the IPF lung. Increased SPP1^hi^ MER Proto-Oncogene, Tyrosine Kinase (MERTK)^hi^ AMs were reported by Morse et al (Morse et al., 2019), who also identified Fatty Acid Binding Protein 4 (FABP4)^hi^ and FCN1^hi^ macrophages. Interestingly, SPP1^hi^ macrophages in IPF and Chronic obstructive pulmonary disease (COPD) appear to have intermediate features between AMs and interstitial macrophages (Adams et al., 2020; Ayaub et al., 2021;(Schupp et al., 2019). Several dendritic cell types were also observed in these scRNAseq studies. Overall, these published findings highlight the diverse nature of myeloid cell subtypes in IPF.

During the recent COVID-19 pandemic, infiltrating monocytes, macrophages and other myeloid cells were identified as key inflammatory cells in patients suffering from severe disease (Bingham et al., 2022; Chen et al., 2022; Sefik et al., 2022). In progressive COVID-19 (also known as long COVID-19) and Post-acute sequelae of COVID (PASC) profibrotic macrophages were identified with a gene signature that mirrored that of profibrotic macrophages identified in IPF (Wendisch et al., 2021). Bosteels C *et al* (Bosteels et al., 2022) confirmed that there is a deficit of alveolar macrophages in PASC-F lungs, perhaps due to defective GM-CSF instruction. At present, the function and contribution of these myeloid cells to the fibrotic process in COVID-19 remains unclear.

Caveolin-1(Cav)-1 is a 20 kDa protein, which is a key protein in the formation of plasma membrane invaginations together with cavin proteins. These plasma membrane invaginations, or caveolae, are present in many cell types in the lung and the reduction in the abundance of caveolae has been shown to contribute to lung diseases including asthma, fibrosis, COPD, acute lung injury and inflammation (Wicher et al., 2019). The Cav-1 protein sequence consists of four domains: (a) NH2-terminal domain, (b) a CSD (residues 82–101) carrying a cholesterol recognition/interaction consensus sequence; (c) a membrane domain, which interacts with membrane lipids; and (d) a COOH-terminal domain (Volonte & Galbiati, 2020). Cav-1 contributes to cell signaling pathways in the normal lung. Decreased expression of Cav-1 is observed in both IPF and PASC-F lung tissue sections and this loss of Cav-1 leads to increased fibroblast activation (Volonte et al., 2002; Xiao et al., 2006). In addition, Cav-1 gene therapy attenuated bleomycin-induced pulmonary fibrosis and reduced infiltration of neutrophils and monocytes/macrophages (Lin et al., 2019). Thus, Cav-1 has an important regulatory role during fibrosis in the lung.

To explore the role of Cav-1 in myeloid cell biology, we examined the effects of LTI-2355 on the functional and synthetic properties of IPF and PASC-F myeloid cells isolated from lung explants. LTI-2355 is a soluble and proteolysis-resistant 13-mer peptide caveolin scaffolding domain (CSD) peptide with anti-fibrotic properties in several pre-clinical models (unpublished). Specifically, we addressed the effect of LTI-2355 on both the functional and synthetic properties on adherent CD45^+^ myeloid cells enriched from normal donor and lung explants from IPF and PASC-F patients.

## Methods and Materials

### CD45^+^ myeloid cell enrichment from normal donor and explanted human lung samples

Single cell solution was obtained from human normal donor and IPF lungs after enzymatic digestion with 10X Liberase (Sigma, 5401127001) and DNaseI (9003-98-9, STEMCELL Technologies) for 40 min at 37°C in complete HBSS (21-023-CV, Corning). The enzymatic digestion was stopped with cold HBSS with bovine serum albumin (BSA), and single cell preparations were obtained by passing the cell mixture through a PluriStrainer (500 – 70µM cell strainers; 43-50500, PluriSelect) before blocking with Fc block for 10 min (422302, BioLegend). CD45^+^ cells were enriched using anti-CD45 magnetic MicroBeads and LS columns (130-045-801 and 130-042-401, Miltenyi Biotec). For CD206-positive (CD206^+^) cell enrichment, cells were incubated for 25 min with biotinylated anti-human MMR (321118, BioLegend), washed, and conjugated with streptavidin microbeads (130-048-102, Miltenyi Biotec) for 15 min. An Institutional Review Board (IRB) at Cedars-Sinai Medical Center approved the use of de-identified human lung samples to enrich for the CD45^+^ myeloid cells studied in the experiments described herein. The normal donor and fibrotic patient characteristics are summarized in **Table 1**.

**Table 1.**
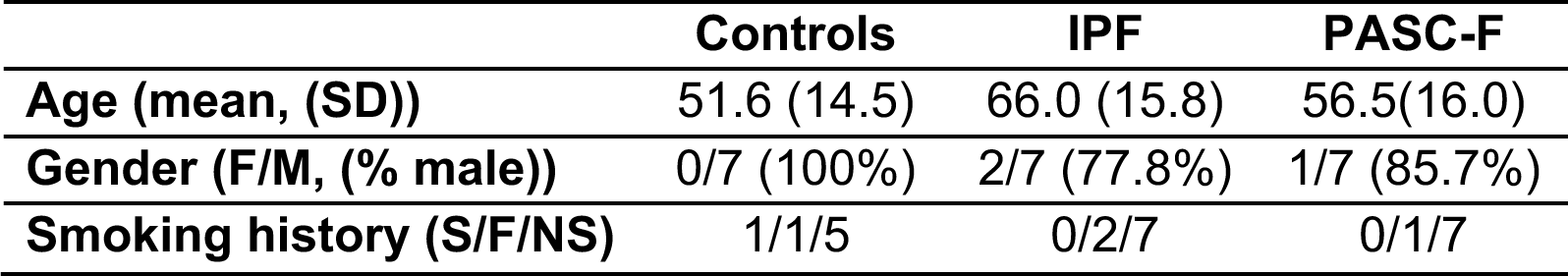
Patient characteristics.

|  | **Controls** | **IPF** | **PASC-F** |
| --- | --- | --- | --- |
| **Age (mean, (SD))** | 51.6 (14.5) | 66.0 (15.8) | 56.5(16.0) |
| **Gender (F/M, (% male))** | 0/7 (100%) | 2/7 (77.8%) | 1/7 (85.7%) |
| **Smoking history (S/F/NS)** | 1/1/5 | 0/2/7 | 0/1/7 |

### CD45^+^ myeloid cell enrichment and *in vitro* cultures

Myeloid cells enriched from normal donor and IPF patient explant lung samples were allowed to adhere for 24 h in Dulbecco’s Modified Eagle Medium (DMEM, 12-604Q Lonza) supplemented with 10% FBS, 1% penicillin-streptomycin-Amphotericin B (17-754E, Lonza), 0.2% Primocin (Invivogen), and 200mM L-glutamine. The non-adherent cellular fraction was removed, and the remaining adherent CD45^+^ myeloid cells were treated with a 10-fold dilution of LTI-2355 in (0.1□M, 1.0□M and 10.0□M), 5 µg/ml UNO peptide (318897, NovoPro, China), or the standard of care drug nintedanib (Ofev®, Boehringer Ingelheim, Germany) at a clinically relevant dose of 80nM.

### Efferocytosis, phagocytosis, and proliferation assays

For quantification of efferocytosis in cultured CD45^+^ myeloid cells from normal donors, IPF, and PASC-F, cellular debris (defined as cellular material (≤ 225□m^2^) was labeled and quantified using IncuCyte 2021 Software in an IncuCyte S3 system. Thirty (30) minutes pre-incubation with 10µM Cytochalasin D (Sigma-Aldrich C8273-1MG) was used as a negative control. For quantification of phagocytosis, CD45^+^ myeloid cells from normal donors, IPF, and PASC-F were cultured with 0.01mg/ml pH-rodo Red Staphylococcus aureus (SA), *E Coli* or zymosan bioparticles (4619/4615/4617, Sartorius, USA) and the Total Red Object Integrated Intensity (Red Calibrated Unit (RCU) x µm²/Image) was measured after compensating for background fluorescence at 1 h intervals using the IncuCyte S3 system (Essen BioScience). Proliferation/viability was quantified in cultured CD45^+^ myeloid cells from normal donors, IPF, and PASC-F with IncuCyte® NucLight Rapid Red Dye (4717, Sartorius). LTI-2355, UNO, nintedanib, or appropriate control substance were added at the indicated concentrations at the start of these assays unless otherwise stated.

### Proteomic analysis

Cell-free tissue culture supernatants was collected after 3 days of culture. Pro-inflammatory and pro-fibrotic mediators were measured in 50µl of cell-free tissue culture supernatants using a 37-Plex Bio-Plex Pro™ Human Inflammation Panel 1 (Bio-Rad #171AL001M) according to manufacturer’s instructions. Briefly, supernatant was incubated for 1 h at 24°C with capture antibodies covalently coupled to magnetic beads. Unbound supernatant was washed before sequential incubation with biotinylated section antibody and streptavidin-phycoerythrin conjugate. Data was acquired using a Bio-Plex 200 reader (Bio Rad). LTI-2355, UNO, nintedanib, or appropriate control substance were added at the indicated concentrations at the start of these assays unless otherwise stated.

## Results

### Aberrant activity and altered morphology of CD45^+^ lung myeloid cells from IPF and PASC-F patients compared with normal donor CD45^+^ myeloid cells

Compared with normal donor myeloid cells (**Figure 1A&B**) at days 0 and 3 in culture, IPF (**Figure 1C&D**) and PASC-F (**Figure 1E&F**) myeloid cells exhibited decreased motility and adherence and showed an altered cell morphology. To examine intrinsic myeloid cell activity, the ability of myeloid cells to clean dead cells and cellular debris (i.e., particulate matter σ; 225□m^2^) was quantified with live imaging during the first 24 h of culture (**Figure 1A-F**). Area under the curve analysis showed impaired clearing of dead cells and particulate debris by IPF myeloid cells when compared to normal donor myeloid cells (p < 0.001, **Figure 1G**).

**Figure 1.**
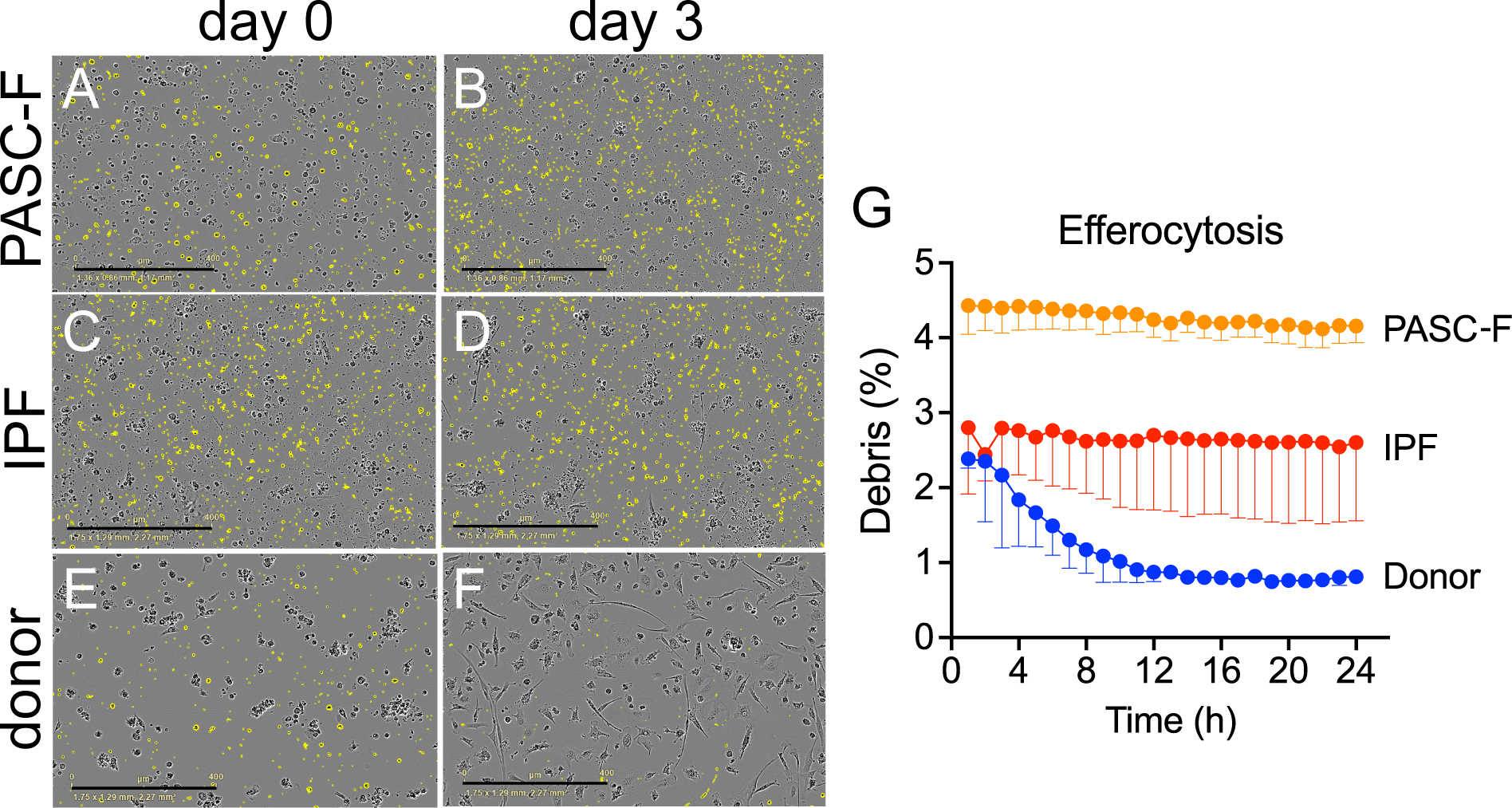
Impaired removal of dead cells and cellular debris by cultured IPF and PASC-F CD45^+^ myeloid cells. CD45^+^ myeloid cells were enriched from normal donor lungs and IPF and PASC-F lung explants and cultured for 3 days with live imaging conducted every hour. Dead cells and cellular debris, defined as particulate material (≤ 225□m^2^), was labeled and quantified using IncuCyte 2021 Software. Representative images of dead cells and cellular debris (colored yellow) in cultures of PASC-F myeloid cells at day 0 (**A**) and 3 (**B**), cultured IPF myeloid cells at day 0 (**C**) and 3 (**D**) and normal donor myeloid cells at day 0 (**E**) and 3 (**F**) of culture. Quantification of debris cleaning by normal donor, IPF, and PASC-F myeloid cells during 24h of culture is summarized in Panel **G**. Data are median with interquartile range and statistical significance was determined using ANOVA and Kruskal-Wallis test, ****p <0.0001.

### Impaired phagocytic activity of CD45^+^ myeloid cells enriched IPF and PASC-F lung explants

To determine whether impaired debris cleaning was due to an impaired phagocytotic response, we added pH-rodo-labelled *Staphylococcus aureus* (SA) beads to isolated myeloid cells. In the presence of pH-rodo SA bioparticles, normal donor myeloid cells showed IPF-derived myeloid cells showed a time-dependent increase in phagocytic index (**Figure 2A** (day 0 in culture), **Figure 2B**, (day 3 in culture), and **Figure 2G**). In contrast, IPF lung myeloid cells exhibited a delayed and impaired intrinsic phagocytic activity in culture (p<0.001, **Figure 2C** (day 0), **Figure 2D** (day 3), and **Figure 2G**). Impaired phagocytic activity was also observed in PASC-F primary myeloid cells (**Figure 2E** (day 0), **Figure 2F** (day 3), and **Figure 2G**). To determine whether uptake of foreign particles was due to an overall impairment of the myeloid cells or specific to SA or a Toll Like Receptor (TLR)-2/4 mechanism, we repeated experiments with *E coli-* and zymosan-coated bioparticles. Coated bioparticle uptake by myeloid cells showed a patient specific but not an agonist coated-bead specific response, indicating an overall intrinsic defect in the function of IPF and PASC-F myeloid cells (**Supplementary Figure 1**). Unlike normal donor myeloid cells in culture, IPF myeloid cells formed aggregates with dead cells and debris and appeared to move around the culture plate with this cellular debris attached to the cell surface (**Supplementary Figure 2**). Decreased synthetic activity in IPF and PASC-F myeloid cells was observed following the analysis of several soluble pro-inflammatory and pro-fibrotic mediators as shown in **Figure 3A**. The one exception among the mediators analyzed was the pro-fibrotic factor chitinase 3-like 1, which was elevated in cultures of both IPF and PASC-F myeloid cells compared with normal donor myeloid cells (**Figure 3A**). Total soluble CD163 (sCD163) and IL-8 levels in cultures of IPF and PASC-F myeloid cells were statically significantly lower when compared to normal donor myeloid cells (**Figure 3B and C**). While secreted osteopontin (OPN) was lower in cultures of IPF and PASC-F myeloid cells (**Figure 3D**), the levels of this cytokine were positively correlated with phagocytic activity of myeloid cells from normal donor and fibrotic lung samples (**Figure 3E**; p = 0.046).

**Figure 2.**
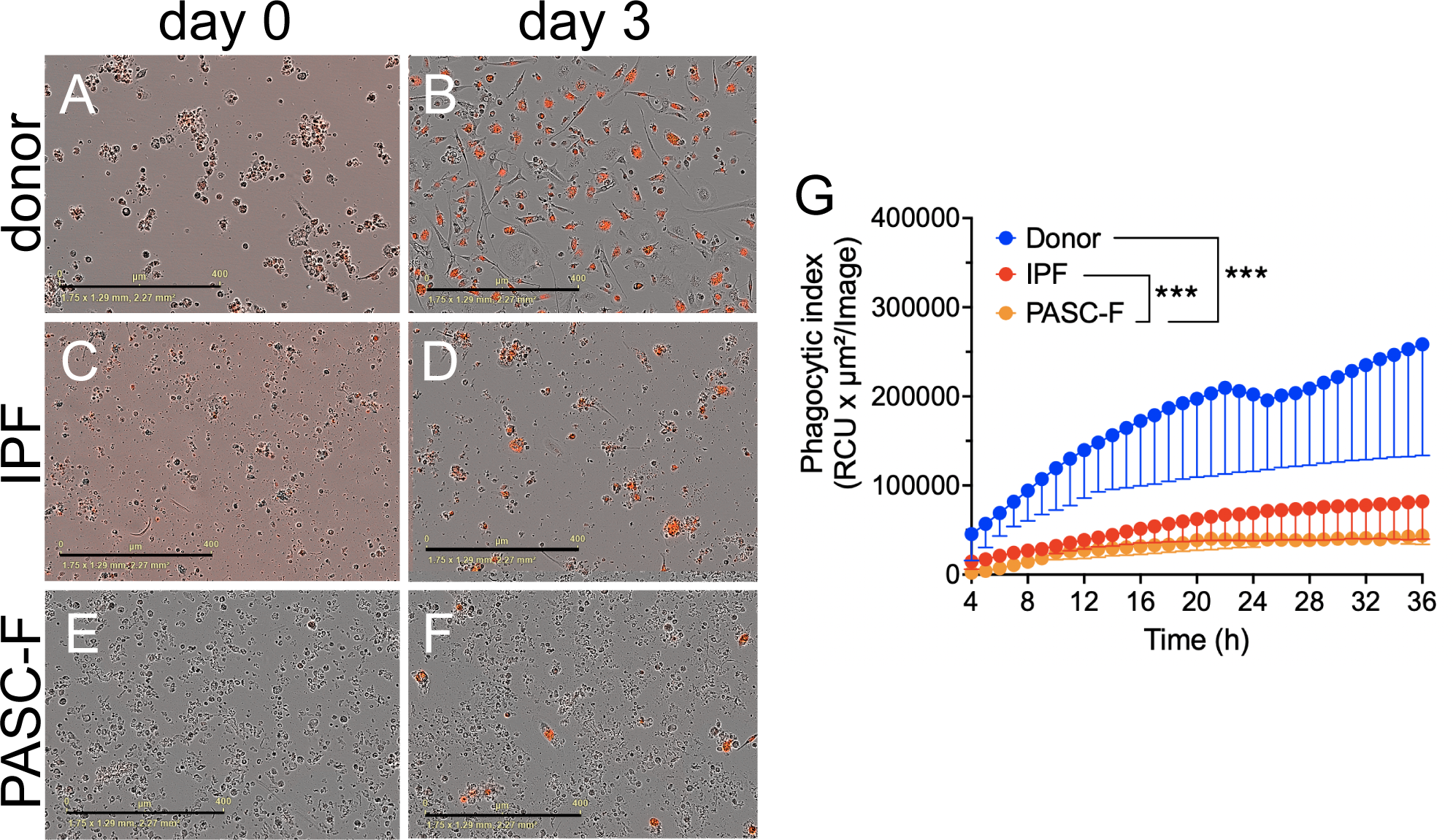
Impaired phagocytosis of bacterial antigen coated bioparticles by IPF and PASC-F myeloid cells. CD45^+^ myeloid cells were enriched from normal donor lungs and IPF and PASC-F lung explants and cultured for 3 days with imaging conducted every hour in the presence of pH-rodo SA beads. Uptake of bioparticles was quantified by measuring the red image fluorescent signal using IncuCyte 2021 Software. Representative images of bioparticle uptake (red label signal) in cultures of PASC-F myeloid cells at day 0 (**A**) and 3 (**B**), cultured IPF myeloid cells at day 0 (**C**) and 3 (**D**) and normal donor myeloid cells at day 0 (**E**) and 3 (**F**) of culture. Quantification of pH-rodo emission by normal donor, IPF, and PASC-F myeloid cells during 24h of culture (**G**). Data are presented as mean ± SEM of three replicates with range, and statistical significance was determined using ANOVA and Kruskal-Wallis test, ***p <0.001.

**Figure 3.**
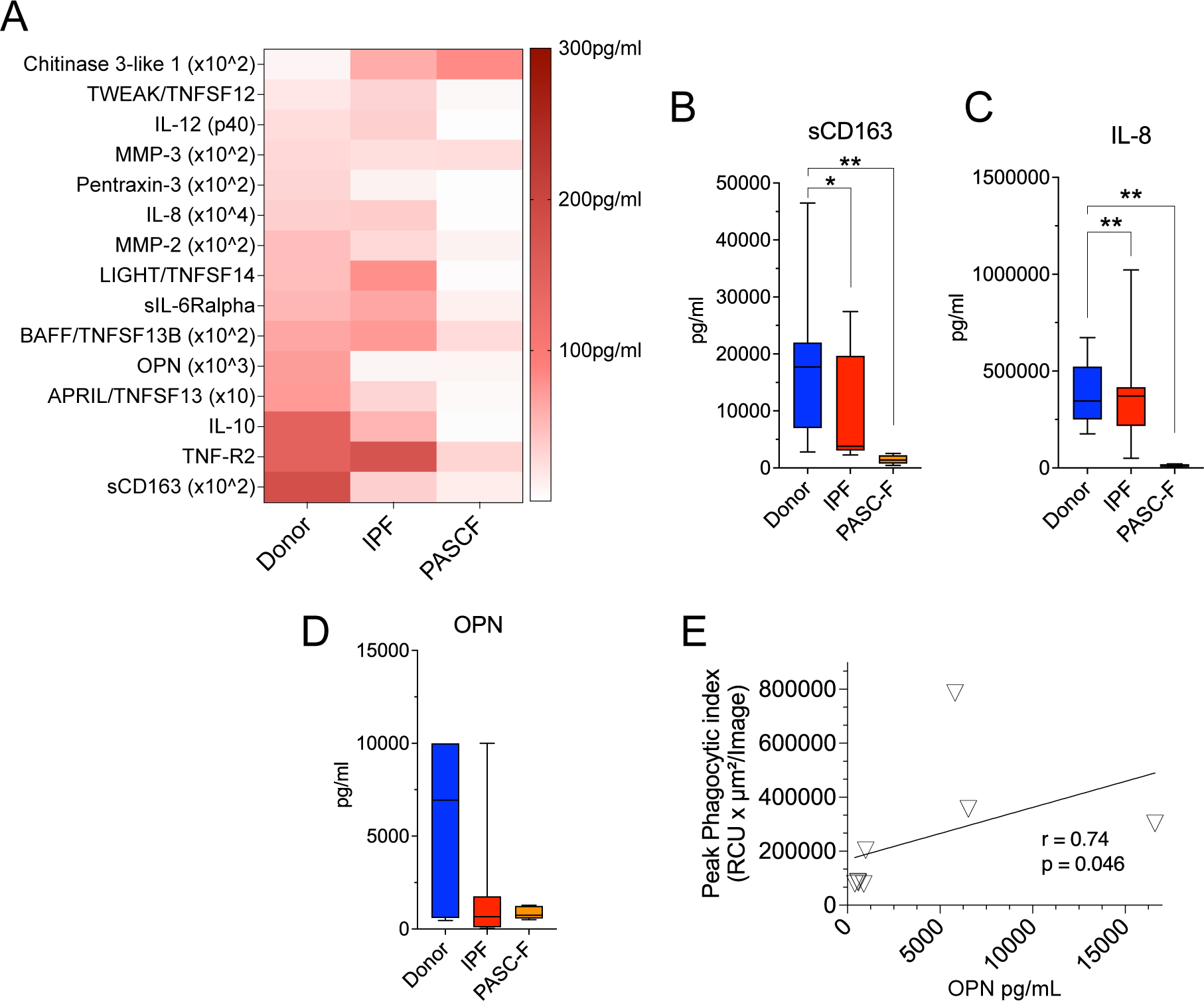
Soluble mediators generated by normal donor, IPF, and PASC-F myeloid cells. CD45^+^ myeloid cells enriched from normal donor, IPF, and PASC-F lung samples or explants were allowed to adhere in tissue culture plates for 24 h. The non-adherent cellular fraction was removed, and the adherent myeloid cells were maintained in culture for an additional 5 days. Protein levels were measured by Bio-Plex (Bio-Rad) in cell-free culture supernatants (**A**). Statistically significant differences in the levels of sCD163 (**B**), IL-8 (**C**), and osteopontin (OPN)(**D**) were observed between normal donor and fibrotic lung myeloid cells. Correlation of OPN levels and peak phagocytic activity by IPF myeloid cells at 48 h of culture is shown in (**E**). Statistical significance was determined using ANOVA and the Mann Whitney U test (**B-D**) and Spearman correlation (**E**); *p <0.05, **p <0.01.

### LTI-2355 increased the phagocytic activity and reduced levels of pro-inflammatory and pro-fibrotic mediators secreted by lung myeloid cells from IPF patients

Normal donor myeloid cells exhibited a time-dependent increase in phagocytic index, which was not enhanced by the presence of LTI-2355 at a dose of 0.1 □M (**Figure 4A**). In contrast, LTI-2355 significantly enhanced the phagocytic activity of IPF myeloid cells at a dose of 1.0□M but not at lower (i.e., 0.1 □M) or higher (i.e., 10 □M) doses (**Figure 4B**). Since increased phagocytic activity of lung myeloid cells is also known to enhance the generation of pro-inflammatory and pro-fibrotic mediators, we examined the synthetic activity of normal donor and IPF myeloid cells. Analysis of supernatants from normal donor (**Figure 4C**) and IPF (**Figure 4D**) myeloid cells showed an overall statistically significant reduction in the presence of several mediators generated by IPF myeloid cells but not normal donor myeloid cells. Specifically, sCD163, IFN-α2, IFN-γ, IL-2, IL-10, IL-12p40, and MMP-1 levels were significantly lower in LTI-2355-treated cultures of IPF myeloid cells compared to control untreated IPF myeloid cells (**Figure 4D**). Interestingly, nintedanib (at 80 nM) did not significantly reduce the levels of any of the soluble mediators measured in cultures of treated IPF myeloid cells versus untreated IPF myeloid cells (**Figure 4D**). It was also interesting to note that there were IPF myeloid cells that did not appear to respond to the presence of a single LTI-2355 treatment after 48 h in culture (**Figure 4E (responders) versus Figure 4F (non-responders**). Analysis of soluble pro-inflammatory mediators in these two groups of IPF patients confirmed that LTI-2355 significantly inhibited the generation of pro-inflammatory and pro-fibrotic mediators in the responder group but not in the non-responder group (**Figure 4G and 4F**, respectively). However, the addition of LTI-2355 every 24 h in culture for the duration of the analysis was found to significantly increase the phagocytic activity of these IPF non-responders (**Supplementary Figure 3**) highlighting the potential of repeated administration of LTI-2355 to IPF myeloid cells as a strategy to enhance the phagocytic properties of these cells.

**Figure 4.**
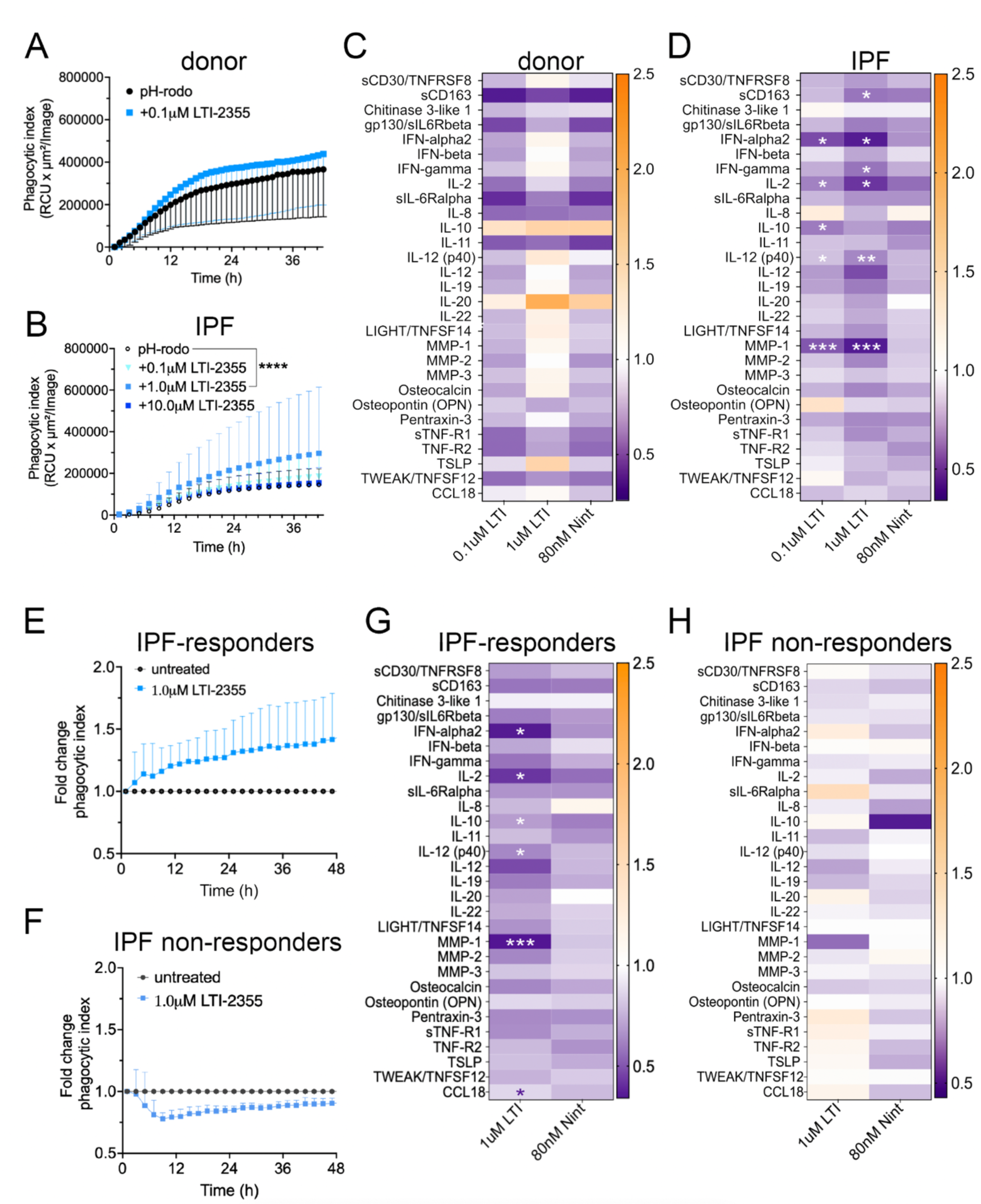
LTI-2355 significantly enhanced the phagocytic activity and decreased soluble mediator levels in enriched CD45+ myeloid cells from IPF patient explants. CD45^+^ myeloid cells were enriched from normal donor lung (**A**) and IPF (**B**) explant tissue and treated with 0.1 – 10.0 □M LTI-2355 for 48 h. To quantify the phagocytic activity, enriched myeloid cells were cultured in the presence of pH-rodo SA beads and imaged every hour in an IncuCyte S3 system. To determine the effect of LTI-2355 and nintedanib on the generation of inflammatory and profibrotic mediators, cell-free culture supernatants were collected from cultured myeloid cells enriched from normal donor (**C**) and IPF lung explant tissue (**D**) and analyzed by Bio-plex (Bio-Rad). During these experiments, IPF patient myeloid cells from 3 patient lung explants did not appear to respond the single LTI-2355 treatment. Consequently, IPF myeloid cells were separated into responders (**E&G**) and non-responders (**F&H**) and the phagocytic index changes (normalized to the pH-rodo control (Fold Change phagocytic index)) and synthetic activity are shown. Statistical significance of the phagocytic index changes was determine using ANOVA and Kruskal-Wallis test, ****p <0.0001. Protein expression was measured by Bio-Plex (Bio-Rad) and presented as median fold change compared to untreated controls. Two-way ANOVA with Bonferroni correction for multiple comparisons compared to control or LTI-2355 (red); *P<0.05, **p<0.01, ***p<0.001.

### LTI-2355 enhanced the phagocytic properties and modulated the synthetic properties of PASC-F myeloid cells

The addition of LTI-2355 to cultured PASC-F myeloid cells dose-dependently increased the phagocytic activity of these cells and maximal PASC-F myeloid phagocytic cell activity was observed at 10 □M of LTI-2355 (**Figure 5A**). Unlike, IPF myeloid cells, the presence of LTI-2355 had a more modest effect on the synthetic activity of pro-inflammatory and pro-fibrotic mediators generated by cultured PASC-F myeloid cells (**Figure 5B**). Together, these data indicated that LTI-2355 significantly altered the phagocytic and more modestly altered the synthetic activity of PASC-F myeloid cells.

**Figure 5.**
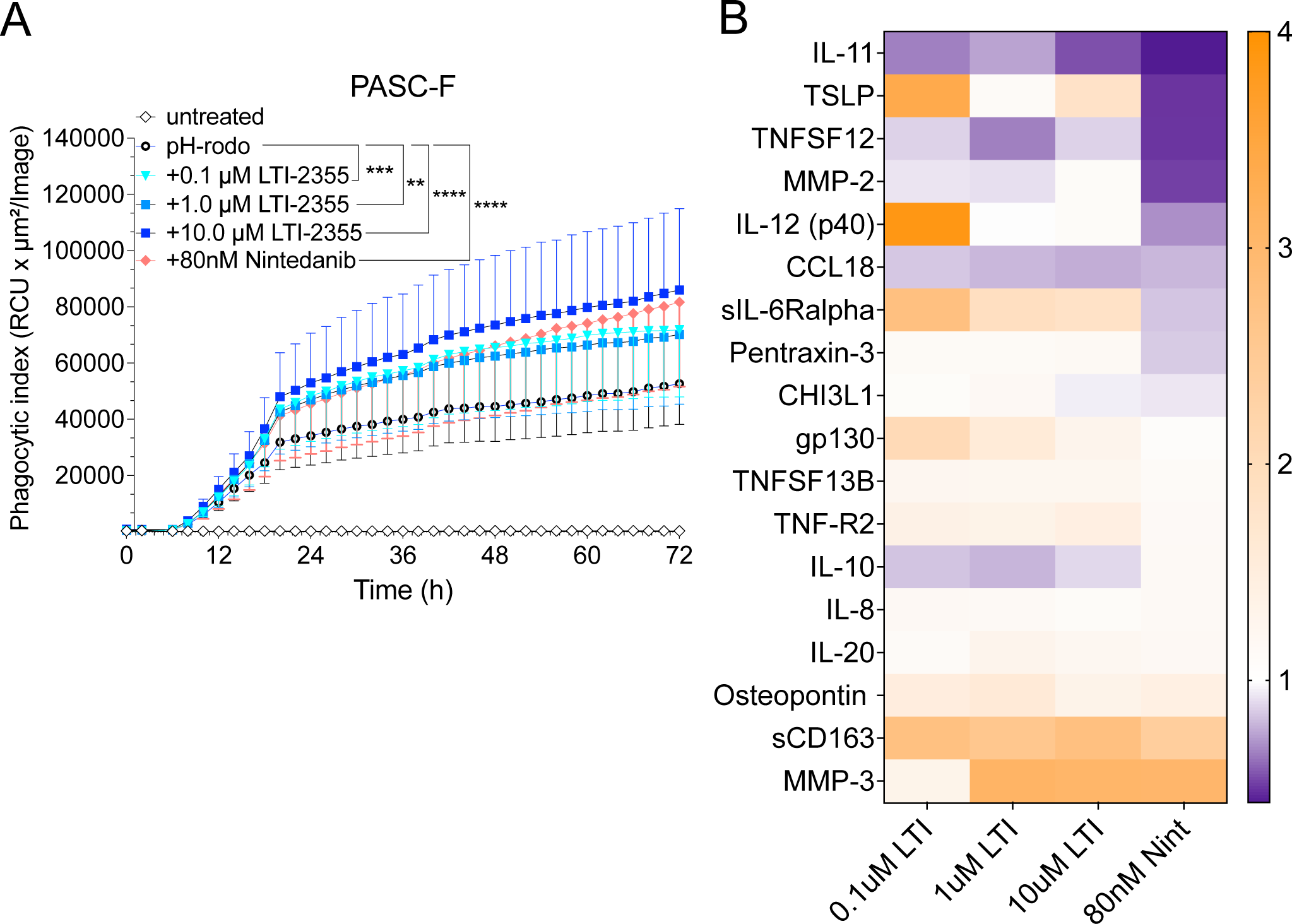
LTI-2355 increased the phagocytic activity of PASC-F myeloid cells. CD45^+^ myeloid cells were enriched from PASC-F patients and treated with 0.1□M, 1□M, 10□M LTI-2355 or Nintedanib (80nM) in culture for 3 days with imaging conducted every hour in presence of pH-rodo SA beads. Uptake of bioparticles was quantified measuring red image fluorescent signal using IncuCyte 2021 Software. Quantification of pH-rodo emission by PASC-F myeloid cells in each treatment group (**A**). Pro-inflammatory protein ratio compared to untreated controls was quantified by Bio-plex in cell culture supernatants of stimulated myeloid cells (B, n = 5). Statistical analysis was determined by ANOVA and Kruskal-Wallis with compared to baseline phagocytic activity (**A**); **p<0.01, ***p<0.001, ****p<0.001 compared with pH-rodo control group.

### Role of CD206 in LTI-2355-mediated effects on IPF and PASC-F myeloid cells

To explore whether CD206 was required for the effects of LTI-2355 on lung myeloid cell phagocytic and synthetic function, we compared the effects of LTI-2355 to those of UNO, which is a CD206 (MRC1) binding peptide (Scodeller et al., 2017). As shown in **Figures 6**, both LTI-2355 and UNO similarly significantly enhanced the phagocytic activity of IPF myeloid cells (**Figure 6A**) and PASC-F myeloid cells (**Figure 6B**). In cultures comprised of CD206-positive myeloid cells, both LTI-2355 and UNO significantly enhances the phagocytic activity of these myeloid cells (**Figure 6C**). In cultures comprised of CD206-negative myeloid cells (from either IPF or PASC-F, the presence of LTI-2355 or nintedanib but not UNO significantly increased the phagocytic activity of this fraction of myeloid cells (**Figure 6D**). Interestingly, nintedanib enhanced the phagocytic activity of CD206-negative but not CD206-positive lung myeloid cells. Also, UNO suppressed the phagocytic activity of CD206-negative lung myeloid cells compared with the pH-rodo control group (**Figure 6D**). LTI2355 and UNO inhibited the synthetic activity of CD206-positive lung myeloid cells as shown in **Figure 6E** but only LTI-2355 showed inhibitory effects on the generation of sCD163 and OPN in CD206-negative lung myeloid cells (**Figure 6F**). Finally, we observed that neither peptide nor nintedanib altered the proliferation of lung myeloid cells from IPF and PASC-F lung explants over 48h in culture (**Supplementary Figure 4**). Overall, these data indicate a broad mechanism of action of LTI-2355 but the presence of CD206 on myeloid cells appears to be required for effects on both phagocytic and synthetic properties of these cells.

**Figure 6.**
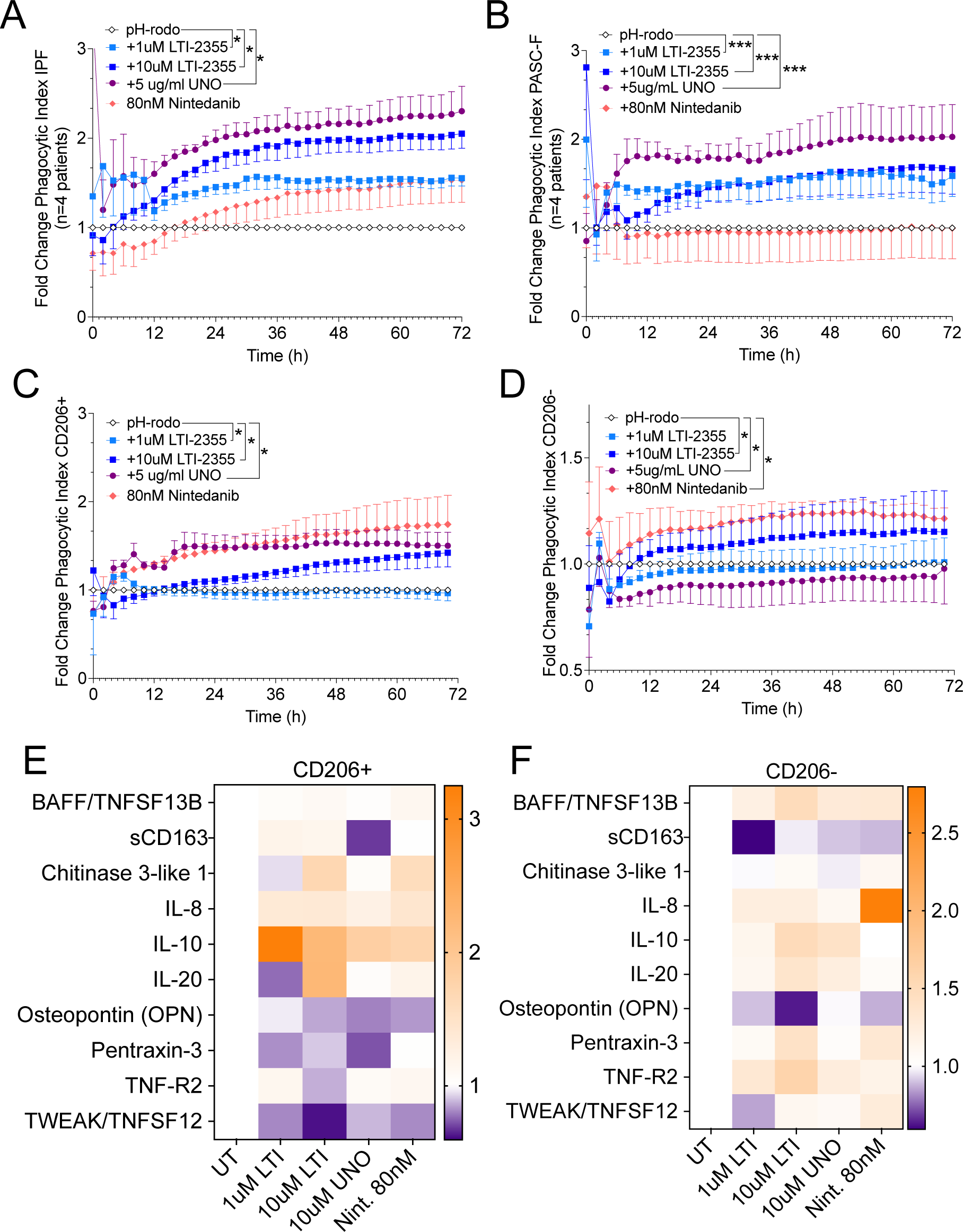
Role of CD206 in the response of IPF and PASC-F myeloid cells to LTI-2355. CD45^+^ myeloid cells were enriched from IPF and PASC-F lung explant tissue and cultured for 3 days with imaging conducted every hour in presence of pH-rodo SA beads. Uptake of bioparticles was quantified measuring red image fluorescent signal using IncuCyte 2021 Software. Quantification of pH-rodo emission by IPF (**A**) and PASC-F (**B**) myeloid cells during 92 h of culture was normalized to the pH-rodo control (Fold Change phagocytic index). Fold Change in the phagocytic index of pH-rodo emission by CD206-positive (CD206^+^) (**C**) and CD206-negative (CD206^-^) (**D**) IPF and PASC-F CD45^+^ enriched myeloid cells normalized to the pH-rodo control. Pro-inflammatory and pro-fibrotic mediator ratios in cell-free tissue culture supernatants from LTI-2355-, UNO- and Nintedanib- (80nM) treated CD206^+^ (**E**) and CD206^-^ (**F**) myeloid cells compared to untreated control groups of both myeloid cell type. Data are presented as mean with standard error of the mean. Multiple comparison with Dunn’s correction, *p <0.05, **p <0.01, ***p <0.001, ***: all conditions vs pH-rodo control (Untreated, UT).

## Discussion

Herein, we demonstrate that CD45^+^ myeloid cells isolated from IPF and PASC-F lung explant tissue have an impaired capacity to clear autologous cellular debris and phagocytose foreign bioparticles in *in vitro* assays. While normal donor myeloid cells efficiently phagocytosed cellular debris and pathogen coated bioparticles, IPF and PASC-F myeloid cells appeared to bind but not engulf cellular debris in the culture system resulting in the clumping of this material on the cell surface of these cells. IPF and PASC-F myeloid cells were also observed traversing around the culture system with this cellular material attached to their cell surface. Uptake of pH-rodo bioparticles was also impaired independently of the nature of the bioparticle highlighting a myeloid cell-intrinsic functional impairment. LTI-2355 dose-dependently improved the efferocytotic and phagocytic properties of both IPF and PASC-F lung myeloid cells while this modified CSD peptide concomitantly reduced the release of pro-inflammatory and pro-fibrotic mediators by these cells. This dual effect of LTI-2355 might be therapeutically significant in the modulation of fibrotic processes in both IPF and PASC-F.

Ours results in clinical fibrotic lung disease confirm findings in mice by Warheit-Niemi HI and colleagues (Warheit-Niemi et al., 2022) who showed that the innate immune response is blunted in bleomycin-exposed mice. Moreover, macrophages loaded with liposomes containing dexamethasone attenuated bleomycin-induced pulmonary fibrosis in mice by reducing the activation of profibrotic macrophages (Ai et al., 2020; Niemann et al., 2021). Mechanistically, exposure to a high burden of apoptotic cells in IPF and PASC-F might have altered the activation state of lung myeloid cells via the downregulation of efferocytosis receptors such as CD11b thereby reducing phagocytic capacity (Kumaran Satyanarayanan et al., 2019; Schif-Zuck et al., 2011). However, in the present study we examined the overall adherent lung myeloid population isolated from human lung explants without distinguishing between satiated (i.e., CD11b^low^ phenotype) or activated (i.e., CD11b^high^ phenotype). Schloesser D *et al* (Schloesser et al., 2023) showed that a suppressed macrophage phenotype was mediated through a cell contact-dependent interaction with senescent fibroblasts. While our results indicate that primary IPF and PASC fibrosis myeloid cells showed an intrinsic suppression of myeloid cell activity, future studies will address the role of the senescent environment on the impairment of IPF and PASC-F lung myeloid cells and whether LTI-2355 alters the functionality of these myeloid cells such that these cells exhibit a pro-resolving phenotype (Zhang et al., 2020).

Studies in murine models have delineated macrophages in an M1 and M2 paradigm, respectively activated by pathogens and Th2 cytokines (Byrne et al., 2015). While M1 macrophages contribute to inflammation by secreting tumor necrosis factor (TNF)-α, interleukin (IL)-1β, IL-6, inducible NO synthase (iNOS) and matrix metalloproteinases (MMPs), M2 macrophages release anti-inflammatory or pro-resolving mediators, including IL-10, transforming growth factor (TGF)-β, arginase (ARG)-1, mannose receptor (CD206), thereby mediating wound repair, tissue remodeling and fibrosis (Murray, 2017; Ogger & Byrne, 2020). CD206 or MRC1 consists of a fibronectin type II domain (FNII) that interacts with collagen, a cysteine-rich domain that binds sulfated proteoglycans, and a lectin domain composed of 8 carbohydrate recognition domains that bind mannose. The importance of the CD206 receptor has been shown in peptide studies. Both Scodeller *et al* and Jaynes *et al* at demonstrated that small peptides, UNO and riptide-182 respectively, are taken up in a CD206-dependent manner into the cell and reprogramed M2-like macrophages to a M1-like macrophage phenotype (Jaynes et al., 2020; Scodeller et al., 2017). The downregulation of cytokine production by LTI-2355 is consistent with a study from Takamura et al who demonstrated that the downregulation of Cav-1 correlated with increased cytokine levels, reversable by Cav-1 transgene administration (Takamura et al., 2019). However, we are cognizant that pro-fibrotic macrophages *in vivo* are not completely identical cell populations compared to M2-like macrophages *in vitro*. Additionally, culture conditions affect macrophage polarity, hence why we chose to perform our experiments on the whole myeloid cell population instead of focusing on M2 macrophages. Furthermore, *in vitro* cultures of macrophages lack alveolar epithelial cells which has been shown to decreases AM phagocytosis and cytokine production in a TGF-β-dependent manner via cell-to-cell contact, nor indirect production of surfactant proteins that can initiate phagocytosis of apoptotic cells and pathogens (Haczku, 2008; Lambrecht, 2006).

In further study, we observed that LTI-2355 had superior effects on phagocytic activity of myeloid compared to nintedanib. Nintedanib has been described to prevent macrophage activation and differentiation towards a M2 phenotype (presumably via the blockade of CSF1R), thereby indirectly preventing fibroblast activation, without affecting M1 markers (Bellamri et al., 2019; Huang et al., 2017; Toda et al., 2018; Ying et al., 2021). The relevance of peptide-based macrophage targeting was confirmed by Ghebremedhin A et al who demonstrated that a mannose receptor targeting peptide RP-832c inhibited M2 macrophage activation and attenuated fibrosis in a bleomycin model of pulmonary fibrosis (Ghebremedhin et al., 2023). As new treatment options emerge in IPF directed at inducing apoptosis in senescent lung cells via a senolytic mechanism (Kasam et al., 2019), it is becoming clear that the restoration of efferocytosis and phagocytosis in lung myeloid cells is needed to remove apoptotic cells from the lung. However, finding the exact mechanisms needed to modulate the myeloid cell activity in this manner requires further investigation in IPF, PASC-F, and other progressive ILDs. Thus, future studies addressing the interplay between myeloid cells, alveolar type 2 epithelial cells, basaloid cells and (myo-)fibroblasts via the establishment of a microenvironment favoring homeostasis is essential in the search for new treatments for IPF.

One limitation to our study is that the myeloid cells were obtained from end-stage IPF and PASC-F explants, which precludes our analysis of inflammatory/immune cells in earlier phases of the disease. In addition, IPF patients were older when compared to normal donors, which might influence the phagocytic and synthetic properties of myeloid cell populations. While a single treatment with LTI-2355 significantly increased phagocytosis in most myeloid cell preparations from IPF and PASC-F patients compared to untreated conditions, LTI-2355 did not restore functional activity levels of IPF and PASC-F myeloid cells to that observed in normal donor myeloid cells. An increased fluorescent signal after LTI-2355 treatment might indicate an increased capacity of active phagocytes to engulf bioparticles, or phenotype switch of the phagocytes in culture towards an active phenotype, but the current studies did not confirm if either or both mechanisms were activated by LTI-2355 treatment. Finally, we did not examine the response of subpopulations of macrophages identified by several groups (Adams et al., 2020; Aran et al., 2019; Morse et al., 2019; Reyfman et al., 2019). This limitation was a consequence of our inability to reproducibly isolate enough viable Mertk^+^, TREM2^+^, and other distinct macrophage subtypes for the functional studies described herein.

Overall, we provide evidence that the CSD peptide LTI-2355 enhances the phagocytic and modulates the synthetic activity of IPF and PASC-F myeloid cells. Combined with previously reported therapeutics effects of CSD on the inhibition of fibroblast activation and the restoration of alveolar epithelial cells, these data indicate the potential of CSDs to modulate myeloid cell activity in IPF and PASC-F. Alternative approaches to enhance/restore anti-fibrotic macrophage function should focus on restoring phagocytic activity. Disruption of the CD47-SIRP to enhance macrophage functions (including phagocytosis, antigen presentation, and ADCC) has shown promise in scleroderma (Lerbs et al., 2020). While macrophages or myeloid cells have been the target in IPF, including Galecto’s galectin-3 inhibitor GB0139, and Hoffmann-La Roche’s zinpentraxin (RPM-151; a recombinant human pentraxin-2), neither of these targeting strategies affected the primary endpoint of slowing Forced Vital Capacity decline in clinical trials. Since adverse effects were observed in the treatment arms of these trials, more research is required to understand the overall role of macrophage/myeloid cell biology in IPF patients and other progressive ILDs. In conclusion, LTI-2355 enhances primary lung myeloid cell functional activity and modulates the pro-inflammatory and pro-fibrotic properties of these immune cells in IPF and PASC-F.

## Supplementary Material

**Supplementary Figure 1.**
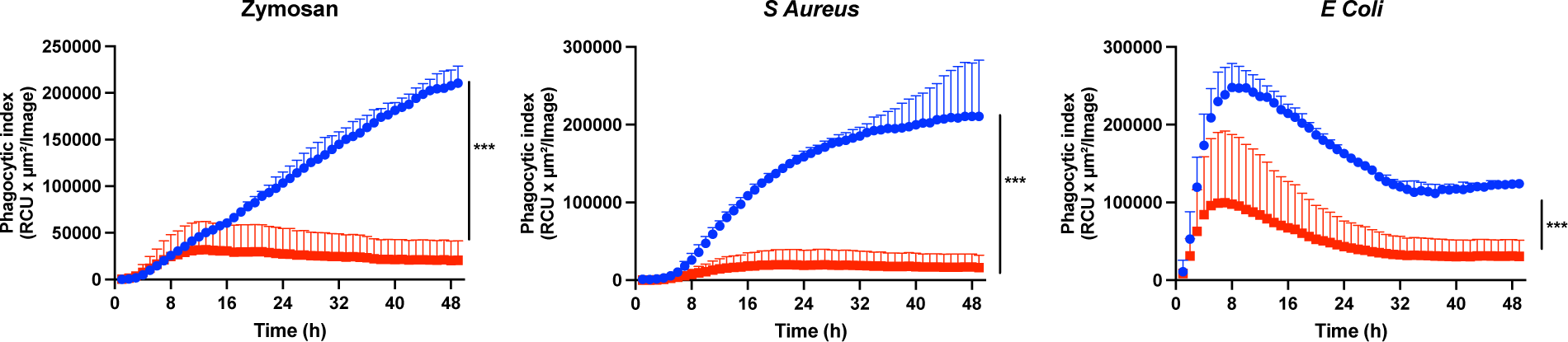
Impaired phagocytosis of bioparticles by IPF myeloid cells was independent of the fungal or bacterial ligands coating the pH-rodo beads. Myeloid cells from IPF lung explant tissue were enriched and cultured for 2 days with imaging every hour in presence of pH-rodo zymosan (A), S Aureus (B) or E Coli (C) beads. Uptake of bioparticles was quantified measuring red image fluorescent signal using IncuCyte 2021 Software. Quantification of pH-rodo emission by control (n=3) and IPF (n=3) myeloid cells during 48h of culture (A). Data are presented as median of three replicates with IQR. Mann Whitney U testing, ****p <0.0001.

**Supplementary Figure 2.**
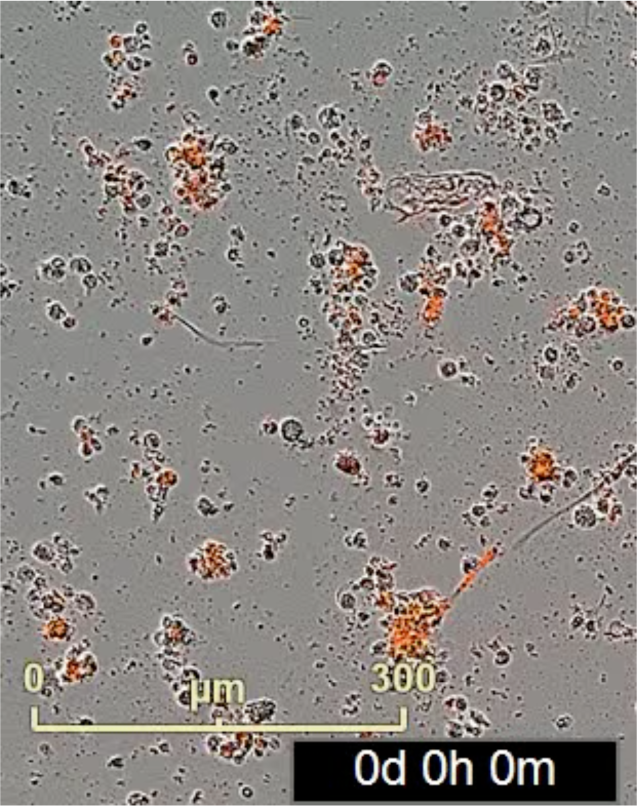
Illustration of aggregate formation and dragging phenotype by IPF myeloid cells. Immune cells from lung explant tissue were enriched and cultured for 3 days. Imaging shown every 6h in presence of pH-rodo SA beads.

**Supplementary Figure 3.**
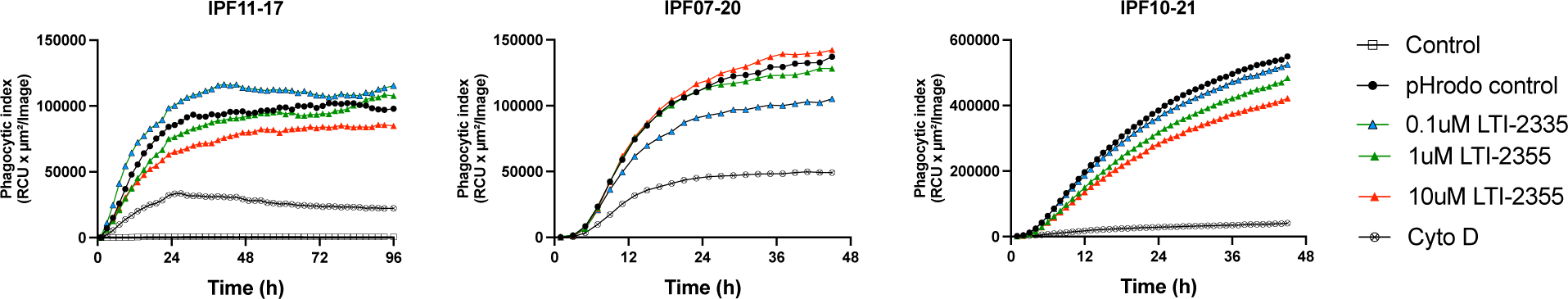
Effect of repeated LTI-2355 administration on the phagocytic index of IPF CD45^+^ myeloid cells. CD45^+^ myeloid cells were enriched from lPF lung explant tissue were enriched and stimulated with either 0.1, 1 or 10 LTI-2355□M at 24 h intervals during this experiment. CD45^+^ myeloid cells from these IPF patients were non-responsive to 1 administration of LTI-2355 into the cultured cells but the repeated administration of this CSD peptide every 24h enhanced the phagocytic of 2 out of the 3 patient myeloid lines examined. The phagocytic index was quantified by live cell imaging using IncuCyte 2021 Software and detected as the red image fluorescent signal resulting from uptake of pH-rodo SA bioparticles.

**Supplementary Figure 4.**
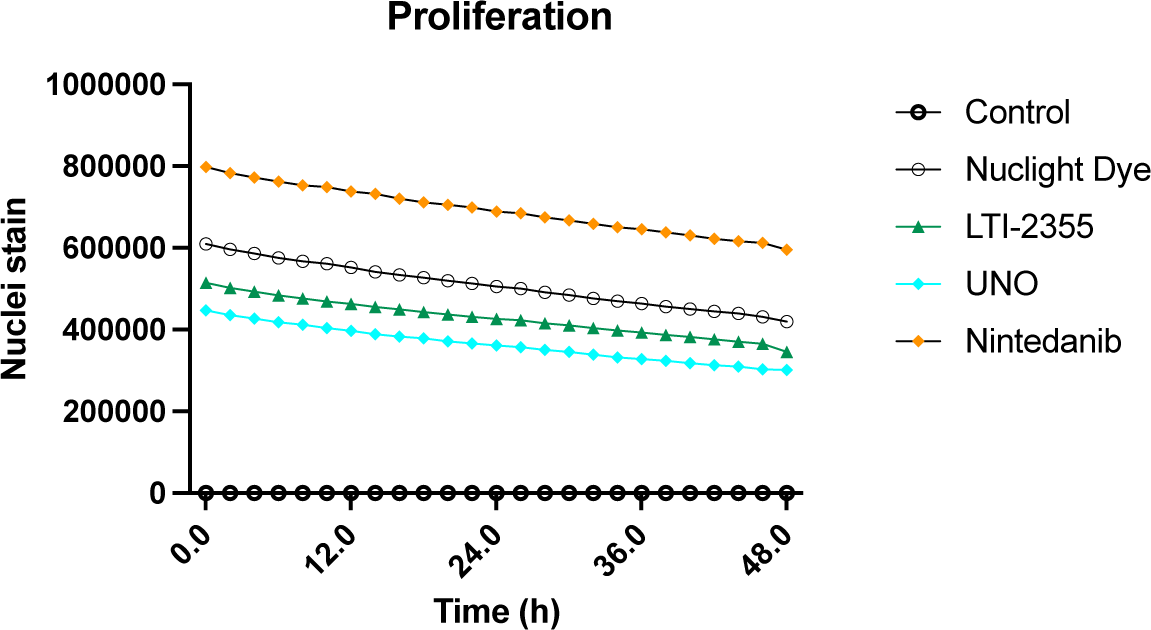
*In vitro* proliferation of IPF myeloid cells was not observed in these cultured cells following exposure to LTI-2355, UNO, or nintedanib. CD45^+^ myeloid cells enriched from IPF lung explant tissue (n=4) were enriched and stimulated after 24h in culture with LTI-2355, Nintedanib or UNO and responses were compared to control medium. Proliferation was quantified with IncuCyte® NucLight Rapid Red Dye (IncuCyte) for 48h with imaging every two hours. Proliferation was quantified by measuring Red Calibrated Unit (RCU) based on the red image fluorescent signal cell-by-cell analysis using IncuCyte 2021 Software.

## Abbreviations

AM: Alveolar macrophage.
CAV: Caveolin
CD: Cluster of differentiation
CH3IL1: Chitinase 3-like-1
COPD: Chronic obstructive pulmonary disease
CSD: Caveolin scaffolding domain
DMEM: Dulbecco’s Modified Eagle Medium
ECM: Extracellular matrix
FABP: Fatty Acid Binding Protein
IL: Interleukin
IPF: Idiopathic Pulmonary Fibrosis
MERTK MER: Proto-Oncogene, Tyrosine Kinase
MMR/MRC: Mannose receptor
MMP: Matrix metalloprotease
Mo-MA: Monocyte derived macrophage
PASC-F: Post-acute sequelae of COVID fibrosis
RCU: Red Calibrated Unit
SA: Staphylococcus aureus
sCD: soluble
CD SPP/OPN: Osteopontin
TLR: Toll like receptor
TNF: Tumor necrosis factor
TR-AM: Tissue-resident alveolar macrophage

## Acknowledgements

This work was supported by National Institute of Health grant P01-HL108793 and Lung Therapeutics, Inc

## Notes

### Competing Interest Statement

BreAnne MacKenzie is an employee of Lung Therapeutics, Inc. Cory Hogaboam is a consulting CSO for Lung Therapeutics, Inc.

### Summary of Updates

This version of the manuscript has been revised to update the funding sources. This work was supported by National Institute of Health grant P01-HL108793 and Lung Therapeutics, Inc

## References

Adams, T. S., Schupp, J. C., Poli, S., Ayaub, E. A., Neumark, N., Ahangari, F., Chu, S. G., Raby, B. A., Deiuliis, G., Januszyk, M., Duan, Q., Arnett, H. A., Siddiqui, A., Washko, G. R., Homer, R., Yan, X., Rosas, I. O., & Kaminski, N. (2020). Single-cell RNA-seq reveals ectopic and aberrant lung-resident cell populations in idiopathic pulmonary fibrosis. In Sci. Adv (Vol. 6). www.ipfcellatlas.com

Ai, F., Zhao, G., Lv, W., Liu, B., & Lin, J. (2020). Dexamethasone induces aberrant macrophage immune function and apoptosis. Oncology Reports, 43(2), 427. 10.3892/OR.2019.7434

Aran, D., Looney, A. P., Liu, L., Wu, E., Fong, V., Hsu, A., Chak, S., Naikawadi, R. P., Wolters, P. J., Abate, A. R., Butte, A. J., & Bhattacharya, M. (2019). Reference-based analysis of lung single-cell sequencing reveals a transitional profibrotic macrophage. Nature Immunology, 20(2), 163–172. 10.1038/S41590-018-0276-Y

Ayaub, E., Poli, S., Ng, J., Adams, T., Schupp, J., Quesada-Arias, L., Poli, F., Cosme, C., Robertson, M., Martinez-Manzano, J., Liang, X., Villalba, J., Lederer, J., Chu, S., Raby, B., Washko, G., Coarfa, C., Perrella, M., El-Chemaly, S., … Rosas, I. (2021). Single Cell RNA-seq and Mass Cytometry Reveals a Novel and a Targetable Population of Macrophages in Idiopathic Pulmonary Fibrosis. BioRxiv, 2021.01.04.425268. 10.1101/2021.01.04.425268

Bellamri, N., Morzadec, C., Joannes, A., Lecureur, V., Wollin, L., Jouneau, S., & Vernhet, L. (2019). Alteration of human macrophage phenotypes by the anti-fibrotic drug nintedanib. International Immunopharmacology, 72, 112–123. 10.1016/J.INTIMP.2019.03.061

Bingham, G. C., Muehling, L. M., Li, C., Huang, Y., Abebayehu, D., Noth, I., Sun, J., Woodfolk, J. A., Barker, T. H., & Bonham, C. (2022). Reduction in circulating monocytes correlates with persistent post-COVID pulmonary fibrosis in multi-omic comparison of long-haul COVID and IPF. 10.1101/2022.09.30.22280468

Bosteels, C., Van Damme, K. F. A., De Leeuw, E., Declercq, J., Maes, B., Bosteels, V., Hoste, L., Naesens, L., Debeuf, N., Deckers, J., Cole, B., Pardons, M., Weiskopf, D., Sette, A., Weygaerde, Y. Vande, Malfait, T., Vandecasteele, S. J., Demedts, I. K., Slabbynck, H., … Lambrecht, B. N. (2022). Loss of GM-CSF-dependent instruction of alveolar macrophages in COVID-19 provides a rationale for inhaled GM-CSF treatment. Cell Reports Medicine, 3(12). 10.1016/j.xcrm.2022.100833

Byrne, A. J., Maher, T. M., & Lloyd, C. M. (2016). Pulmonary Macrophages: A New Therapeutic Pathway in Fibrosing Lung Disease? Trends in Molecular Medicine, 22(4), 303–316. 10.1016/J.MOLMED.2016.02.004

Byrne, A. J., Mathie, S. A., Gregory, L. G., & Lloyd, C. M. (2015). Pulmonary macrophages: key players in the innate defence of the airways. Thorax, 70(12), 1189–1196. 10.1136/THORAXJNL-2015-207020

Chen, S. T., Park, M. D., del Valle, D. M., Buckup, M., Tabachnikova, A., Thompson, R. C., Simons, N. W., Mouskas, K., Lee, B., Geanon, D., D’Souza, D., Dawson, T., Marvin, R., Nie, K., Zhao, Z., LeBerichel, J., Chang, C., Jamal, H., Akturk, G., … Merad, M. (2022). A shift in lung macrophage composition is associated with COVID-19 severity and recovery. Science Translational Medicine, 14(662), 5168. 10.1126/SCITRANSLMED.ABN5168/SUPPL_FILE/SCITRANSLMED.ABN5168_MDAR_REPRODUCIBILITY_CHECKLIST.PDF

Ding, L., Zeng, Q., Wu, J., Li, D., Wang, H., Lu, W., Jiang, Z., & Xu, G. (2017). Caveolin-1 regulates oxidative stress-induced senescence in nucleus pulposus cells primarily via the p53/p21 signaling pathway in vitro. Molecular Medicine Reports, 16(6), 9521. 10.3892/MMR.2017.7789

Ghebremedhin, A., Salam, A. Bin, Adu-Addai, B., Noonan, S., Stratton, R., Ahmed, M. S. U., Khantwal, C., Martin, G. R., Lin, H., Andrews, C., Karanam, B., Rudloff, U., Lopez, H., Jaynes, J., & Yates, C. (2023). A Novel CD206 Targeting Peptide Inhibits Bleomycin-Induced Pulmonary Fibrosis in Mice. Cells, 12(9), 1254. 10.3390/CELLS12091254/S1

Gu, L., Surolia, R., Larson-Casey, J. L., He, C., Davis, D., Kang, J., Antony, V. B., & Carter, A. B. (2021). Targeting Cpt1a-Bcl-2 interaction modulates apoptosis resistance and fibrotic remodeling. Cell Death & Differentiation 2021 29:1, 29(1), 118–132. 10.1038/s41418-021-00840-w

Haczku, A. (2008). Protective role of the lung collectins surfactant protein A and surfactant protein D in airway inflammation. The Journal of Allergy and Clinical Immunology, 122(5), 861–879. 10.1016/J.JACI.2008.10.014

Huang, J., Maier, C., Zhang, Y., Soare, A., Dees, C., Beyer, C., Harre, U., Chen, C. W., Distler, O., Schett, G., Wollin, L., & Distler, J. H. W. (2017). Nintedanib inhibits macrophage activation and ameliorates vascular and fibrotic manifestations in the Fra2 mouse model of systemic sclerosis. Annals of the Rheumatic Diseases, 76(11), 1941– 1948. 10.1136/annrheumdis-2016-210823

Inoshima, I., Kuwano, K., Hamada, N., Yoshimi, M., Maeyama, T., Hagimoto, N., Nakanishi, Y., & Hara, N. (2004). Induction of CDK inhibitor p21 gene as a new therapeutic strategy against pulmonary fibrosis. American Journal of Physiology. Lung Cellular and Molecular Physiology, 286(4). 10.1152/AJPLUNG.00209.2003

Jaynes, J. M., Sable, R., Ronzetti, M., Bautista, W., Knotts, Z., Abisoye-Ogunniyan, A., Li, D., Calvo, R., Dashnyam, M., Singh, A., Guerin, T., White, J., Ravichandran, S., Kumar, P., Talsania, K., Chen, V., Ghebremedhin, A., Karanam, B., bin Salam, A., … Rudloff, U. (2020). Mannose receptor (CD206) activation in tumor-associated macrophages enhances adaptive and innate antitumor immune responses. In Sci. Transl. Med (Vol. 12). www.imperial.ac.uk/research/animallectins/

Kasam, R. K., Reddy, G. B., Jegga, A. G., & Madala, S. K. (2019). Dysregulation of mesenchymal cell survival pathways in severe fibrotic lung disease: The effect of nintedanib therapy. Frontiers in Pharmacology, 10(MAY), 532. 10.3389/FPHAR.2019.00532/BIBTEX

Kiss, A. L., & Botos, E. (2009). Endocytosis via caveolae: alternative pathway with distinct cellular compartments to avoid lysosomal degradation? Journal of Cellular and Molecular Medicine, 13(7), 1228–1237. 10.1111/J.1582-4934.2009.00754.X

Kumaran Satyanarayanan, S., el Kebir, D., Soboh, S., Butenko, S., Sekheri, M., Saadi, J., Peled, N., Assi, S., Othman, A., Schif-Zuck, S., Feuermann, Y., Barkan, D., Sher, N., Filep, J. G., & Ariel, A. (2019). IFN-β is a macrophage-derived effector cytokine facilitating the resolution of bacterial inflammation. Nature Communications, 10(1). 10.1038/S41467-019-10903-9

Lambrecht, B. N. (2006). Alveolar macrophage in the driver’s seat. Immunity, 24(4), 366–368. 10.1016/J.IMMUNI.2006.03.008

Larson-Casey, J. L., Deshane, J. S., Ryan, A. J., Thannickal, V. J., & Carter, A. B. (2016). Macrophage Akt1 Kinase-Mediated Mitophagy Modulates Apoptosis Resistance and Pulmonary Fibrosis. Immunity, 44(3), 582–596. 10.1016/J.IMMUNI.2016.01.001

Lerbs, T., Cui, L., King, M. E., Chai, T., Muscat, C., Chung, L., Brown, R., Rieger, K., Shibata, T., & Wernig, G. (2020). CD47 prevents the elimination of diseased fibroblasts in scleroderma. JCI Insight, 5(16). 10.1172/JCI.INSIGHT.140458

Lin, X., Barravecchia, M., Matthew Kottmann, R., Sime, P., & Dean, D. A. (2019). Caveolin-1 gene therapy inhibits inflammasome activation to protect from bleomycin-induced pulmonary fibrosis. Scientific Reports, 9(1). 10.1038/s41598-019-55819-y

Marudamuthu, A. S., Bhandary, Y. P., Fan, L., Radhakrishnan, V., MacKenzie, B. A., Maier, E., Shetty, S. K., Nagaraja, M. R., Gopu, V., Tiwari, N., Zhang, Y., Watts, A. B., Williams, R. O., Criner, G. J., Bolla, S., Marchetti, N., Idell, S., & Shetty, S. (2019). Caveolin-1-derived peptide limits development of pulmonary fibrosis. Science Translational Medicine, 11(522). 10.1126/SCITRANSLMED.AAT2848

McCarron, A., Cmielewski, P., Drysdale, V., Parsons, D., & Donnelley, M. (2022). Effective viral-mediated lung gene therapy: is airway surface preparation necessary? Gene Therapy 2022, 1–9. 10.1038/s41434-022-00332-7

Mergia, A. (2017). The Role of Caveolin 1 in HIV Infection and Pathogenesis. Viruses, 9(6). 10.3390/V9060129

Misharin, A. v., Morales-Nebreda, L., Reyfman, P. A., Cuda, C. M., Walter, J. M., McQuattie-Pimentel, A. C., Chen, C. I., Anekalla, K. R., Joshi, N., Williams, K. J. N., Abdala-Valencia, H., Yacoub, T. J., Chi, M., Chiu, S., Gonzalez-Gonzalez, F. J., Gates, K., Lam, A. P., Nicholson, T. T., Homan, P. J., … Perlman, H. (2017). Monocyte-derived alveolar macrophages drive lung fibrosis and persist in the lung over the life span. Journal of Experimental Medicine, 214(8), 2387–2404. 10.1084/jem.20162152

Morse, C., Tabib, T., Sembrat, J., Buschur, K. L., Bittar, H. T., Valenzi, E., Jiang, Y., Kass, D. J., Gibson, K., Chen, W., Mora, A., Benos, P. v., Rojas, M., & Lafyatis, R. (2019). Proliferating SPP1/MERTK-expressing macrophages in idiopathic pulmonary fibrosis. European Respiratory Journal, 54(2). 10.1183/13993003.02441-2018

Mundy, D. I., Machleidt, T., Ying, Y. S., Anderson, R. G. W., & Bloom, G. S. (2002). Dual control of caveolar membrane traffic by microtubules and the actin cytoskeleton. Journal of Cell Science, 115(Pt 22), 4327–4339. 10.1242/JCS.00117

Murray, P. J. (2017). Macrophage Polarization. Annual Review of Physiology, 79, 541–566. 10.1146/ANNUREV-PHYSIOL-022516-034339

Niemann, S., Lucarini, L., Mae Gowdy, K., Yang, J., Sang, X., Wang, Y., Xue, Z., Qi, D., Fan, G., Tian, F., & Zhu, Y. (2021). Macrophage-Targeted Lung Delivery of Dexamethasone Improves Pulmonary Fibrosis Therapy via Regulating the Immune Microenvironment. Frontiers in Immunology | Www.Frontiersin.Org, 12, 613907. 10.3389/fimmu.2021.613907

Ogger, P. P., & Byrne, A. J. (2020). Macrophage metabolic reprogramming during chronic lung disease. Mucosal Immunology 2020 14:2, 14(2), 282–295. 10.1038/s41385-020-00356-5

Povedano, J. M., Martinez, P., Serrano, R., Tejera, Á., Gómez-López, G., Bobadilla, M., Flores, J. M., Bosch, F., & Blasco, M. A. (2018). Therapeutic effects of telomerase in mice with pulmonary fibrosis induced by damage to the lungs and short telomeres. ELife, 7. 10.7554/ELIFE.31299

Reyfman, P. A., Walter, J. M., Joshi, N., Anekalla, K. R., McQuattie-Pimentel, A. C., Chiu, S., Fernandez, R., Akbarpour, M., Chen, C. I., Ren, Z., Verma, R., Abdala-Valencia, H., Nam, K., Chi, M., Han, S. H., Gonzalez-Gonzalez, F. J., Soberanes, S., Watanabe, S., Williams, K. J. N., … Misharin, A. v. (2019). Single-cell transcriptomic analysis of human lung provides insights into the pathobiology of pulmonary fibrosis. American Journal of Respiratory and Critical Care Medicine, 199(12), 1517–1536. 10.1164/rccm.201712-2410OC

Ruigrok, M. J. R., Frijlink, H. W., Melgert, B. N., Olinga, P., & Hinrichs, W. L. J. (2021). Gene therapy strategies for idiopathic pulmonary fibrosis: recent advances, current challenges, and future directions. 10.1016/j.omtm.2021.01.003

Schif-Zuck, S., Gross, N., Assi, S., Rostoker, R., Serhan, C. N., & Ariel, A. (2011). Saturated-efferocytosis generates pro-resolving CD11b low macrophages: modulation by resolvins and glucocorticoids. European Journal of Immunology, 41(2), 366–379. 10.1002/EJI.201040801

Schloesser, D., Lindenthal, L., Sauer, J., Chung, K. J., Chavakis, T., Griesser, E., Baskaran, P., Maier-Habelsberger, U., Fundel-Clemens, K., Schlotthauer, I., Watson, C. K., Swee, L. K., Igney, F., Park, J. E., Huber-Lang, M. S., Thomas, M. J., El Kasmi, K. C., & Murray, P. J. (2023). Senescent cells suppress macrophage-mediated corpse removal via upregulation of the CD47-QPCT/L axis. The Journal of Cell Biology, 222(2). 10.1083/JCB.202207097/VIDEO-1

Schupp, J. C., Adams, T., Neumark, N., Poli De Frias, S., Ahangari, F., Deiuliis, G., Chu, S., Yan, X., Kaminski, N., Prasse, A., & Rosas, I. O. (2019). Macrophage Programs in BAL and Lung Parenchyma of the Healthy and in IPF Patients. www.atsjournals.org

Scodeller, P., Simón-Gracia, L., Kopanchuk, S., Tobi, A., Kilk, K., Säälik, P., Kurm, K., Squadrito, M. L., Kotamraju, V. R., Rinken, A., de Palma, M., Ruoslahti, E., & Teesalu, T. (2017). Precision Targeting of Tumor Macrophages with a CD206 Binding Peptide. Scientific Reports, 7(1). 10.1038/s41598-017-14709-x

Sefik, E., Qu, R., Junqueira, C., Kaffe, E., Mirza, H., Zhao, J., Brewer, J. R., Han, A., Steach, H. R., Israelow, B., Blackburn, H. N., Velazquez, S. E., Chen, Y. G., Halene, S., Iwasaki, A., Meffre, E., Nussenzweig, M., Lieberman, J., Wilen, C. B., … Flavell, R. A. (2022). Inflammasome activation in infected macrophages drives COVID-19 pathology. Nature 2022 606:7914, 606(7914), 585–593. 10.1038/s41586-022-04802-1

Shetty, S., Marudamuthu, A., Bhandary, Y., Fan, L., Radhakrishnan, V., MacKenzie, B., Maier, E., Shetty, S. K., Gopu, V., Tiwari, N., Zhang, Y., Watts, A. B., Williams, R. O. I., Idell, S., Criner, G. J., Bolla, S., Marchetti, N., & Nagaraja, M. R. (2020). Caveolin-1-Derived Peptide Limits Development of Pulmonary Fibrosis. ATS 2020 International Conference American Thoracic Society International Conference Meetings Abstracts, A7881–A7881. 10.1164/AJRCCM-CONFERENCE.2020.201.1_MEETINGABSTRACTS.A7881

Takamura, N., Yamaguchi, Y., Watanabe, Y., Asami, M., Komitsu, N., & Aihara, M. (2019). Downregulated Caveolin-1 expression in circulating monocytes may contribute to the pathogenesis of psoriasis. Scientific Reports 2019 9:1, 9(1), 1–12. 10.1038/s41598-018-36767-5

Toda, M., Mizuguchi, S., Minamiyama, Y., Yamamoto-Oka, H., Aota, T., Kubo, S., Nishiyama, N., Shibata, T., & Takemura, S. (2018). Pirfenidone suppresses polarization to M2 phenotype macrophages and the fibrogenic activity of rat lung fibroblasts. Journal of Clinical Biochemistry and Nutrition, 63(1), 58. 10.3164/JCBN.17-111

Volonte, D., & Galbiati, F. (2020). Caveolin-1, a master regulator of cellular senescence. Cancer Metastasis Reviews, 39(2), 397–414. 10.1007/S10555-020-09875-W

Volonte, D., Zhang, K., Lisanti, M. P., & Galbiati, F. (2002). Expression of caveolin-1 induces premature cellular senescence in primary cultures of murine fibroblasts. Molecular Biology of the Cell, 13(7), 2502–2517. 10.1091/MBC.01-11-0529

Warheit-Niemi, H. I., Edwards, S. J., SenGupta, S., Parent, C. A., Zhou, X., O’Dwyer, D. N., & Moore, B. B. (2022). Fibrotic lung disease inhibits immune responses to staphylococcal pneumonia via impaired neutrophil and macrophage function. JCI Insight, 7(4). 10.1172/jci.insight.152690

Wendisch, D., Dietrich, O., Mari, T., von Stillfried, S., Ibarra, I. L., Mittermaier, M., Mache, C., Chua, R. L., Knoll, R., Timm, S., Brumhard, S., Krammer, T., Zauber, H., Hiller, A. L., Pascual-Reguant, A., Mothes, R., Bülow, R. D., Schulze, J., Leipold, A. M., … Sander, L. E. (2021). SARS-CoV-2 infection triggers profibrotic macrophage responses and lung fibrosis. Cell, 184(26), 6243–6261.e27. 10.1016/j.cell.2021.11.033

Wicher, S. A., Prakash, Y. S., & Pabelick, C. M. (2019). Caveolae, caveolin-1 and lung diseases of aging. Expert Review of Respiratory Medicine, 13(3), 291–300. 10.1080/17476348.2019.1575733

Xiao, M. W., Zhang, Y., Hong, P. K., Zhou, Z., Feghali-Bostwick, C. A., Liu, F., Ifedigbo, E., Xu, X., Oury, T. D., Kaminski, N., & Choi, A. M. K. (2006). Caveolin-1: a critical regulator of lung fibrosis in idiopathic pulmonary fibrosis. The Journal of Experimental Medicine, 203(13), 2895. 10.1084/JEM.20061536

Yamaguchi, Y., Yasuoka, H., Stolz, D. B., & Feghali-Bostwick, C. A. (2011). Decreased caveolin-1 levels contribute to fibrosis and deposition of extracellular IGFBP-5. Journal of Cellular and Molecular Medicine, 15(4), 957–969. 10.1111/J.1582-4934.2010.01063.X

Ying, H., Fang, M., Hang, Q. Q., Chen, Y., Qian, X., & Chen, M. (2021). Pirfenidone modulates macrophage polarization and ameliorates radiation-induced lung fibrosis by inhibiting the TGF-β1/Smad3 pathway. Journal of Cellular and Molecular Medicine, 25(18), 8662–8675. 10.1111/jcmm.16821

Zhang, F., Ayaub, E. A., Wang, B., Puchulu-Campanella, E., Li, Y.-H., Hettiarachchi, S. U., Lindeman, S. D., Luo, Q., Rout, S., Srinivasarao, M., Cox, A., Tsoyi, K., Nickerson-Nutter, C., Rosas, I. O., & Low, P. S. (2020). Reprogramming of profibrotic macrophages for treatment of bleomycin-induced pulmonary fibrosis. EMBO Molecular Medicine, 12(8), e12034. 10.15252/EMMM.202012034

Zhu, X. D., Zhuang, Y., Ben, J. J., Qian, L. L., Huang, H. P., Bai, H., Sha, J. H., He, Z. G., & Chen, Q. (2011). Caveolae-dependent Endocytosis Is Required for Class A Macrophage Scavenger Receptor-mediated Apoptosis in Macrophages. The Journal of Biological Chemistry, 286(10), 8231. 10.1074/JBC.M110.145888

